# Microscopic and mesoscopic effects of reward uncertainty in monkey fronto-parietal areas

**DOI:** 10.1101/2019.12.17.879262

**Authors:** Bahareh Taghizadeh, Nicholas C. Foley, Saeed Karimimehr, Michael Cohanpour, Mulugeta Semework, Sameer A. Sheth, Reza Lashgari, Jacqueline Gottlieb

**Affiliations:** Brain Engineering Research Center, Institute for Research and Fundamental Sciences, Tehran 19395-5746, Iran; School of Cognitive Sciences, Institute for Research in Fundamental Sciences, Tehran 19395-5746, Iran; Department of Neuroscience, Columbia University; Zuckerman Mind Brain Behavior Institute, Columbia University; Department of Neurosurgery, Baylor College of Medicine; The Kavli Institute for Brain Science, Columbia University

**Author notes:** **Corresponding author:** Jacqueline Gottlieb, PhD, Department of Neuroscience, Columbia University, 3227 Broadway, New York, NY 10025. Equal contribution (Bahareh Taghizadeh, Nicholas C. Foley). **Author Contributions**: NCF and JG designed the experiment. NCF, SAS, MC, MS, RL and JG implemented the experiment and collected the data. BT, SK, RL and JG analyzed the data and wrote the manuscript.

## Abstract

Theories of executive function propose that controlled information processing is costly and is allocated according to the behavioral benefits it brings. Computational theories predict that the benefits of new information depend on prior uncertainty, but the cellular effects of uncertainty on the executive network are incompletely understood. Using simultaneous recordings in monkeys, we reveal several mechanisms by which the fronto-parietal network reacts to uncertainty independently of average reward gains. We show that the variance of expected rewards, independently of the value of the rewards, was represented in single neuron and population spiking activity and local field potential (LFP) oscillations. Moreover, uncertainty asymmetrically affected the coherence between spikes and LFPs, selectively suppressing information transmission from the frontal to the parietal lobe but enhancing transmission from the parietal to the frontal lobe, consistent with Bayesian principles of optimal inference under uncertainty.

## Introduction

Executive control is broadly understood as the ability to engage in information processing in pursuit of a goal, especially in circumstances requiring non-habitual or novel responses^1^. In humans and monkeys, executive function depends on a network of frontal and parietal areas, which is consistently activated in relation to demanding behaviors requiring the suppression of inappropriate response tendencies, monitoring and adjusting behavioral strategies, and the goal-directed control of attention^1, 2^.

Current views of executive function propose that controlled (rather than automatic) information processing is costly and is engaged in proportion to the benefits it brings to the organism^1, 3^. An important implication of this view, in light of Bayesian and predictive coding theories, is that control should be optimally allocated to tasks that not merely have reward value but, more specifically, have uncertainty. It is in conditions of higher *ex ante* uncertainty that animals can expect to obtain the greatest gains in prediction accuracy from processing new information^4–7^.

Consistent with this view, a growing literature shows that attention is recruited by uncertainty independently of reward gains. Animals are intrinsically motivated to resolve uncertainty independently of instrumental incentives^5, 8^ and the intrinsic value of information is encoded in overt (saccadic) attention in humans^9^ and monkeys^10^. Moreover, oculomotor neurons in monkey parietal cortex have stronger responses preceding saccades that are expected to reduce uncertainty relative to those that merely confirm prior expectations, and this sensitivity to uncertainty is independent of reward gains^11^.

And yet, while existing studies have tested neural activity in the fronto-parietal network in tasks involving risk and ambiguity, learning, exploration, novelty, or surprise(e.g., ^12–17^), critical open questions remain about the cellular effects of uncertainty on this network.

One question concerns the distinction between uncertainty and reward gains. In instrumental conditions, when animals make reward-maximizing decisions, reductions of decision uncertainty by definition produce increases in long-term reward gains^5, 18^. While a handful of studies has used non-instrumental conditions to show that individual neurons have distinct responses to the variance and value of expected rewards, these studies have targeted the orbitofrontal cortex^19^ and subcortical structures^20, 21^. In contrast, studies of the fronto-parietal network have yet to examine the encoding uncertainty independent of decision incentives^22–25^.

A second key question is how uncertainty affects not only the neural activity within areas but information flow between areas. It is often proposed that, in states of high prior uncertainty, the brain downregulates top-down signals conveying uncertain prior expectations and upregulates the bottom-up transmission of reliable sensory cues. This uncertainty-dependent weighting is a cornerstone of Bayesian^4^ and predictive coding theories^6^ but there has been no empirical demonstration of uncertainty-dependent modulations of functional connectivity.

To examine these questions, we simultaneously recorded single-neuron responses and local field potential (LFP) oscillations in the dorsolateral prefrontal cortex (dlPFC) and area 7A, two interconnected nodes of the monkey fronto-parietal network. We used a simple task in which monkeys were cued to expect certain or uncertain rewards but could not make decisions to maximize those rewards, allowing us to examine the effects of uncertainty independently of decision incentives. We show that uncertainty has distinct representations in action potential activity and LFP oscillations and asymmetrically enhances spike-field coherence (SFC) from the parietal and frontal lobe while suppressing SFC in the opposite direction, suggesting a modulation of information flow consistent with theoretical predictions of optimal inference under uncertainty.

## Results

### Task and behavior

Two monkeys performed a visually guided saccade task in which they formed expectations about the trial’s rewards based on familiar visual cues. On each trial after achieving central fixation, the monkeys were shown a reward cue at 8° eccentricity to the right or left of fixation, followed by a 400 ms delay period and presentation of the target for the subsequent saccade (**Fig. 1A**). Upon making the required saccade, the monkeys received the reward according to the probability signaled by the cue. Because the saccade direction was instructed, not chosen, and the cue and target locations were independently randomized, this design allowed us to examine reward expectations independently of saccade planning or reward-maximizing decision strategies.

**Fig. 1.**
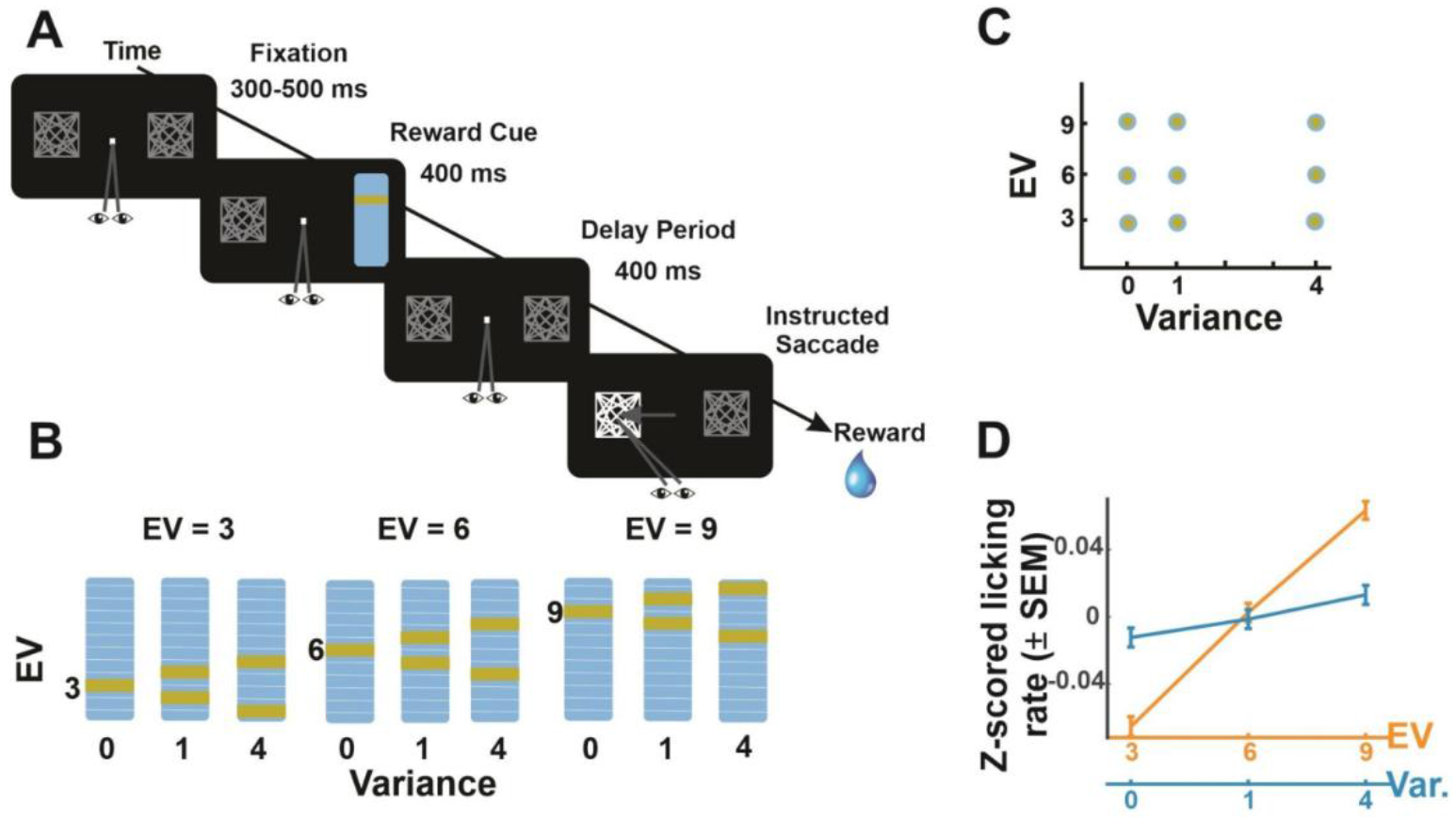
Reward distributions. **(A) Task** The monkeys initiated a trial by looking at a fixation point flanked by two placeholders. A randomly selected placeholder was then replaced by a reward cue for 400 ms, followed by a 400 ms delay (memory) period, and presentation of the saccade target (luminance increase in a randomly selected placeholder). After making the instructed saccade, the monkeys received a reward according to the cued distribution. **(B) Visual cues** Monkeys were familiarized with 9 visual cues signaling the possible reward magnitudes on an 11-point scale (1 point = 0.1 mL of water). **(C) Orthogonalization of variance and EV** Across the cue set, variance and EV could each take 3 discrete levels and were statistically dissociated. **(D) Anticipatory licking** before reward delivery increased as a function of both variance and EV. Points show average and SEM of the z-scored licking rates (n = 12,029 trials).

Monkeys were familiarized with a set of 9 cues signaling 9 reward distributions whose variance and EV were statistically dissociated. Three cues signaled deterministic rewards of, respectively, 3, 6 or 9 points (**Fig. 1B**). The remaining 6 cues indicated probabilistic rewards consisting of a small or large magnitude that were equally likely to occur and were symmetrically positioned, with low or high variance, around the same levels of EV (**Fig. 1B**). This mean-preserving procedure produced a cue set that statistically dissociated 3 levels of variance (0, 1 and 4) and 3 levels of EV (**Fig. 1C**).

Analyses of anticipatory licking confirmed that the monkeys were familiar with the cues and were cognizant of both variance and EV (**Figure 1D**). The GLM coefficients (*Methods*) measuring the modulations of licking were significantly greater than zero for both variance and EV (mean ± SEM across sessions, variance: 0.03±0.0006; EV: 0.24±0.002, all p <10^−9^ relative to 0, signed-rank test). These effects were independent of the location of the visual cue (included in the GLM model as a nuisance regressor) and did not impact the monkeys’ saccades, showing that reward variance and EV invigorated anticipatory licking independently of visual and saccade orienting.

To investigate the neural correlates of variance and EV, we implanted multi-channel electrode arrays in area 7A and the dlPFC focusing on subdivisions that are reciprocally connected and have visual and attention-related activity – i.e., area OPT in the parietal cortex and the pre-arcuate portion of the dlPFC^26, 27^ (**Fig. S1**). We describe the effects of variance and EV in single-neuron activity, followed by the effects on LFP oscillatory power and spike-field coherence (SFC).

### Variance and EV have distinct single-neuron representations

In both 7A and dlPFC, individual neurons showed significant encoding of variance or EV (**Table 1)** but the two response types were clearly segregated (**Fig. 2**). GLM analysis of delay period firing rates (*Methods*) showed that the coefficients of the two factors were uncorrelated (7A: r = 0.03, p = 0.49, n = 522; dlPFC: r = 0.06, p = 0.15; n = 530) and the fraction of neurons with main effects of both factors was below 3% in each area - lower than would be expected by chance.

**Table 1.** Single-neuron sensitivity to variance and EV in dlPFC and 7A. The left half of the table (“Proportion significant”), shows the percentage (number) of neurons with significant coefficients, and the results of chi-square tests of proportions comparing 7A and dlPFC. The right half (“β values) shows the mean ± SEM of the signed coefficients in the sensitive cells and the results of Kruskall-Wallis non-parametric analysis of variance comparing the two areas. Note that, while the individual area averages refer to the *signed* values of the coefficients, the comparison between areas is on the *absolute* values to indicate whether the effects are stronger in any one area regardless of sign

| | Proportion significant | | | $\beta$ values (mean ± SEM) | | |
| --- | --- | --- | --- | --- | --- | --- |
|  | dIPFC | 7A | dIPFC vs 7A | dIPFC | 7A | dIPFC vs 7A |
| **EV** |  |  |  |  |  |  |
| EV+ | 5.7% (30) | 12% (62) | $\chi^2 = 2.410$, df =1, $p > 0.1$ | $1.13 \pm 0.13^*$ | $1.32 \pm 0.11^*$ | $p = 0.72$ |
| EV- | 5.7% (30) | 6% (31) | $\chi^2 = 0.007$, df =1, $p > 0.9$ | $-0.94 \pm 0.07^*$ | $-0.80 \pm 0.07^*$ | $p = 0.06$ |
| All EV sensitive | 11.3% (60) | 18% (93) | $\chi^2 = 1.690$, df =1, $p > 0.1$ | $0.10 \pm 0.15$ | $0.61 \pm 0.13^*$ | $p = 0.70$ |
| **Risk** |  |  |  |  |  |  |
| Variance+ | 2.8% (15) | 2.3% (12) | $\chi^2 = 0.060$, df =1, $p > 0.1$ | $0.73 \pm 0.09^*$ | $0.79 \pm 0.1^*$ | $p = 0.62$ |
| Variance- | 3.4% (18) | 3.1% (16) | $\chi^2 = 0.017$, df =1, $p > 0.9$ | $-0.72 \pm 0.08^*$ | $-0.95 \pm 0.1^*$ | $p = 0.10$ |
| All var. sensitive | 6.2% (33) | 5.4% (28) | $\chi^2 = 0.070$, df =1, $p > 0.1$ | $-0.06 \pm 0.14$ | $-0.20 \pm 0.18$ | $p = 0.08$ |
| **Interaction** |  |  |  |  |  |  |
| All interaction | 6.6% (35) | 6.1% (32) | $\chi^2 = 0.018$, df =1, $p > 0.9$ | $0.04 \pm 0.18$ | $0.82 \pm 0.23^*$ | $p = 0.04$ |
| **All modulations** | 22% (117) | 26% (134) | $\chi^2 = 3.460$, df = 7, $p > 0.1$ | | | |

**Fig. 2.**
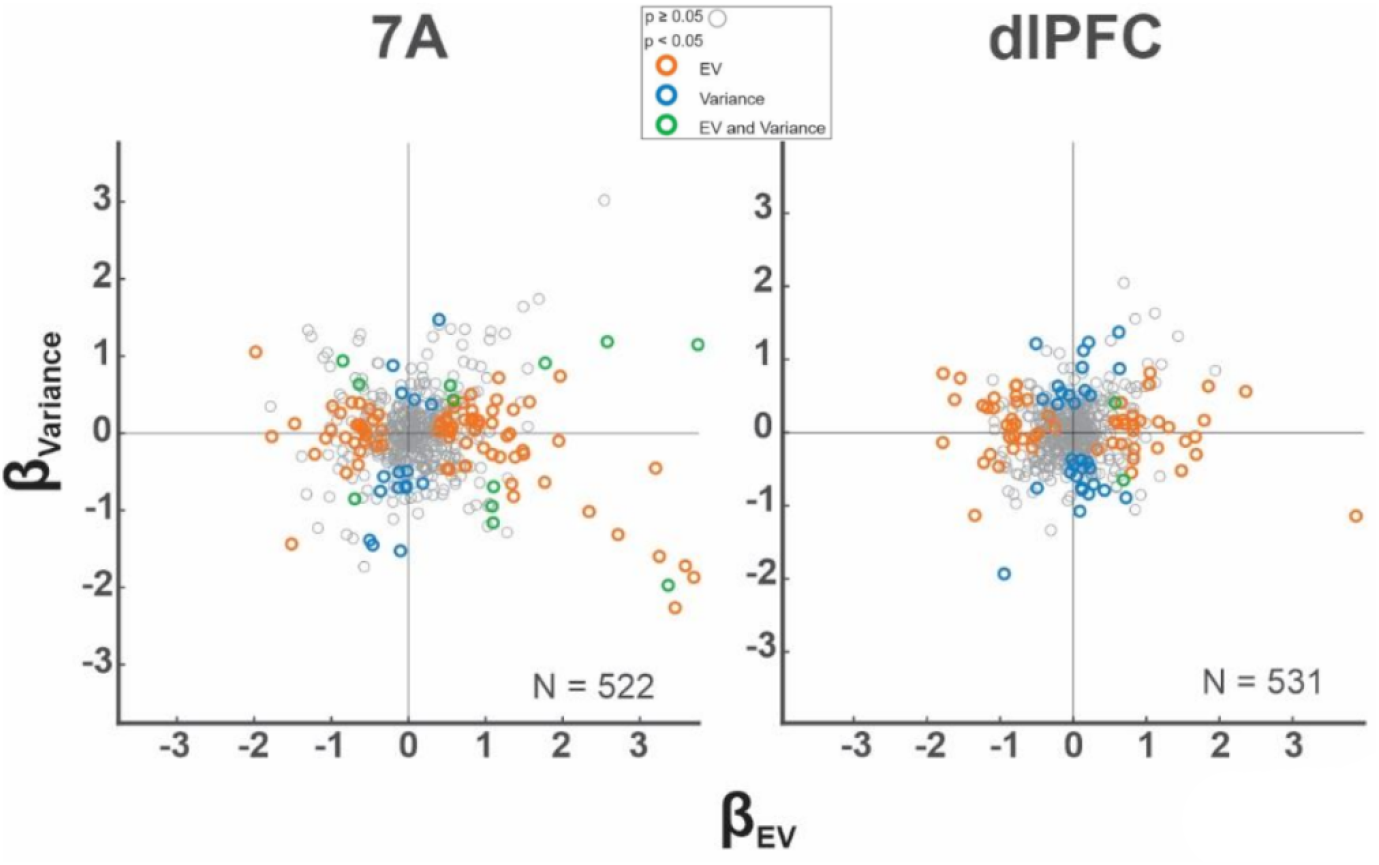
Variance and EV responses are uncorrelated in individual cells. Scatterplots of the GLM coefficients capturing the effect of EV (β_EV_, abscissa) and variance (β_Variance_, ordinate) in individual neurons in 7A and dlPFC. Every point is one neuron, and the colors indicate the significance of the coefficients.

Rather than producing up- or down-modulation of overall firing rates, both uncertainty and EV produced increases or decreases in firing in distinct populations of cells (**Fig. 2**). Responses with positive and negative scaling had similar prevalence and strength in the two areas **(Table 1**). With the single exception of a net positive effect of EV in area 7A (mean ± SEM GLM coefficient of 0.171 ± 0.032, p = 10^−6^ relative to 0) the average coefficients showed no net enhancement or suppression of firing with either variance or EV (all p > 0.23). Subgroups of cells with positive and negative scaling had sustained effects throughout the cue and delay epochs (**Fig. S2A**) and, as noted above, were exclusively selective to one factor (**Fig. S2B**). Although some neurons in both areas showed selectivity for the location of the visual cue as expected, EV and variance-related responses were uncorrelated with visuo-spatial selectivity (**Fig. S3**).

#### Noise correlations

Given the stark segregation of response types we found in both areas, we asked whether different response types were associated with distinct connectivity. To examine this question, we computed noise correlations between trial by trial activity in pairs of simultaneously recorded cells, focusing on firing rates in a 600 ms pre-cue epoch preceding cue onset to avoid confounds related to evoked activity^28^.

Noise correlations were higher in pairs in which both neurons coded for the same factor (both neurons encoding variance or both encoding EV) relative to pairs with mixed selectivity (**Fig. 3A**, “across-factor” vs “within factor”) and this difference was highly robust in both areas (**Table 2,** dlPFC, p = 2.5×10^−7^, n = 47 and n = 56 pairs; 7A p = 5.1×10^−8^, n = 79 and n = 43 pairs; Kruskal-Wallis test). In addition, in pairs with homogeneous selectivity, noise correlations were larger if the two neurons had the same versus opposite polarity (**Fig. 3A**) for both variables and both areas (**Table 2**). Variance, EV or response polarity had no effect on across-trial variability (Fano factor), ruling out that this may have produced apparent effect on noise correlations. Thus, subject to their encoding polarity, neurons responding to variance shared distinct variability relative to those encoding EV.

**Table 2.**
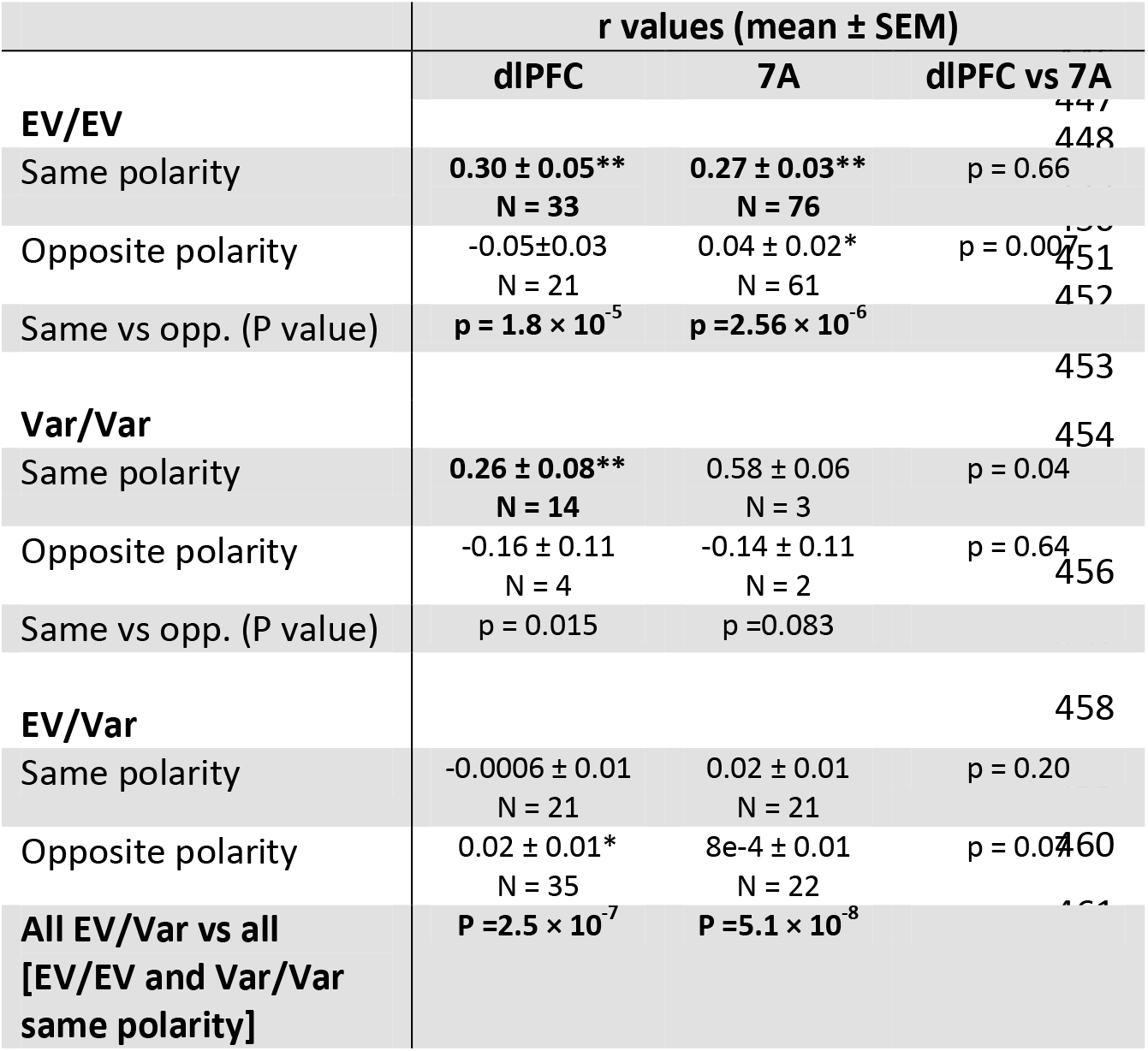
Each entry shows the average and SEM of the Pearson correlation coefficient, and the number (N) of simultaneously recorded cell pairs of each type that met the analysis criteria (see *Methods*). Stars and bold typeface indicate the results of signed-rank tests relative to 0: **\*\*p < 0.01**; *p < 0.05

**Fig. 3.**
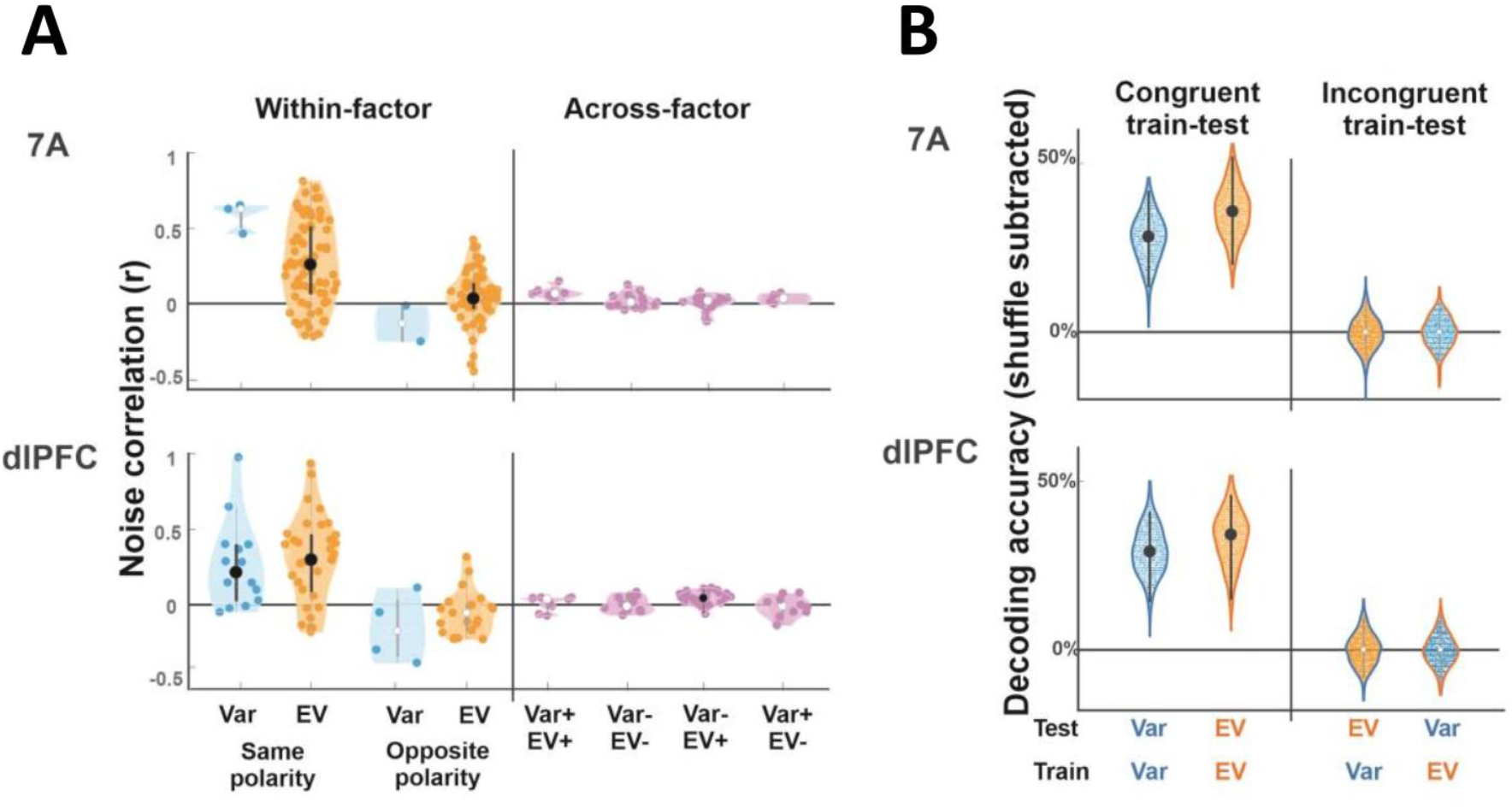
Evidence for separate representations in noise correlations and population decoding. **(A) Noise correlations** Each violin plot shows the distributions of noise correlation coefficients for pairs of cells that were simultaneously recorded and had specific combinations of selectivity as noted on the x-axis. The dots in each distribution show individual pairs; the larger points and whiskers show the median coefficient and 25^th^ and 75^th^ percentiles. Distributions that are significantly higher than 0 (p < 0.05) are shown with black dots and whiskers, otherwise they are shown with white dots and gray whiskers. “Within-factor” distributions show pairs in which both cells were selective to EV or both were selective for variance, further separated by whether the two cells had the same encoding polarity (EV+/EV+ or EV−;/EV−;, variance+/variance+ or variance-/variance-) or opposite polarity (EV+/EV−; or variance+/variance-). “Across-factor” distributions show the coefficients for pairs in which one cell was selective for EV and the other for variance, further separated by polarity as noted. **(B) Classification accuracy based on population responses** Each violin plot shows the distribution of accuracy across 200 bootstrap iterations (after subtracting the accuracy in a randomized dataset). The large dot and error bars show the average accuracy and 95% confidence intervals, with above-chance classification shown with black dots and whiskers. The different distributions correspond to different train/test regimes, as indicated by the x-axis and colors (test variable: dot color; train variable: outline color; orange: EV; blue: variance).

#### Decoding

Because information can be transmitted by neurons that lack linear selectivity, we conducted a final analysis to estimate the decoding capacity from the entire population of cells. We trained support vector machine (SVM) classifiers to perform pairwise discriminations between the different levels of variance and EV based on the population responses and analyzed the boostrapped distributions of excess accuracy (the differences in accuracy in the real and label-shuffled (null) data sets). To determine the extent to which variance and EV had distinct or overlapping representations, we also tested incongruent train-testing regimes – training the classifiers on variance and EV and testing on the untrained variable.

Decoding performance in congruent training-testing regimes was clearly superior to that in incongruent regimes for both variables in both areas. In both the pooled analyses (**Fig. 3B)** and pairwise comparisons **(Fig. S4),**95% confidence bands were clearly above 0 for all congruent train-testing classifications, while decoding in incongruent regimes was significantly weaker and at chance levels in all cases. There were no significant differences between the decoding of variance and EV in 7A and dlPFC.

In sum, analysis of single-neuron activity, noise correlation and population decoding show that variance and EV had clearly segregated representations that were similar in 7A and the dlPFC.

### Variance and EV modulate oscillatory LFP power

Because, in addition to spiking activity, oscillatory LFP potentials are sensitive indicators of cognitive states^29–31^ we next examined how oscillations are affected by variance and EV. To this end, we divided single-trial LFP traces into 1 Hz x 1 ms pixels spanning the cue and delay epochs and, for each pixel, fit a GLM model that included variance and EV as factors, controlling for cue location and interactions (identical to the model applied to spiking activity; *Methods*). The resulting coefficient maps showed that variance and EV exerted consistent effects in two frequency bands: a lower frequency band between 8-18 Hz, corresponding to α/low-β frequencies, and a higher band of 18-43 Hz, corresponding to the high-β/low-β frequencies (**Fig. 4, 5**).

**Fig. 4.**
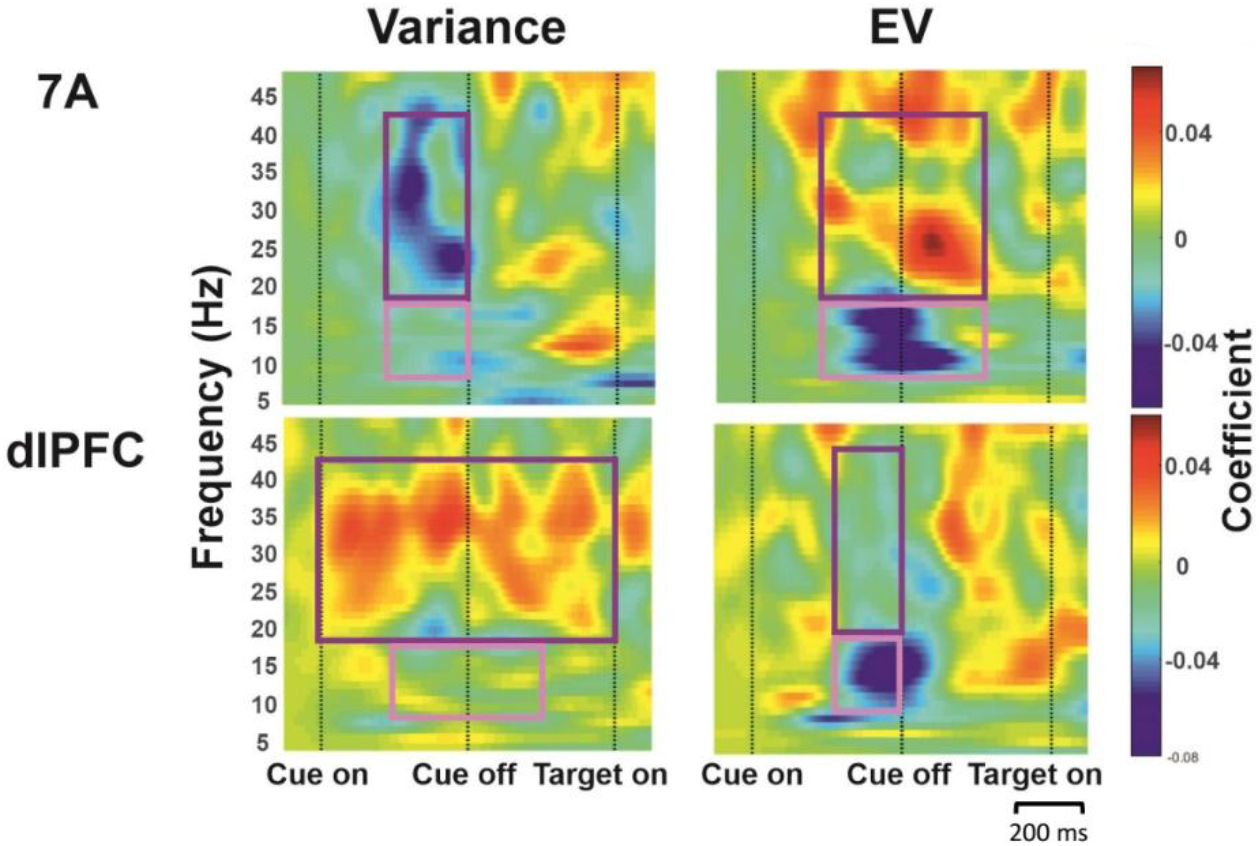
Variance and EV have common and divergent effects on LFP oscillations. Time-frequency maps of GLM coefficients indicating the effects of variance and EV on LFP power aligned on cue onset. The rectangles superimposed on each map indicate the regions of interest (ROI) where consistent effects were found in both monkeys, in the α/low-β (8-18 Hz, pink) and high-β/low-Υ (18-43 Hz, purple) frequency bands.

**Fig. 5.**
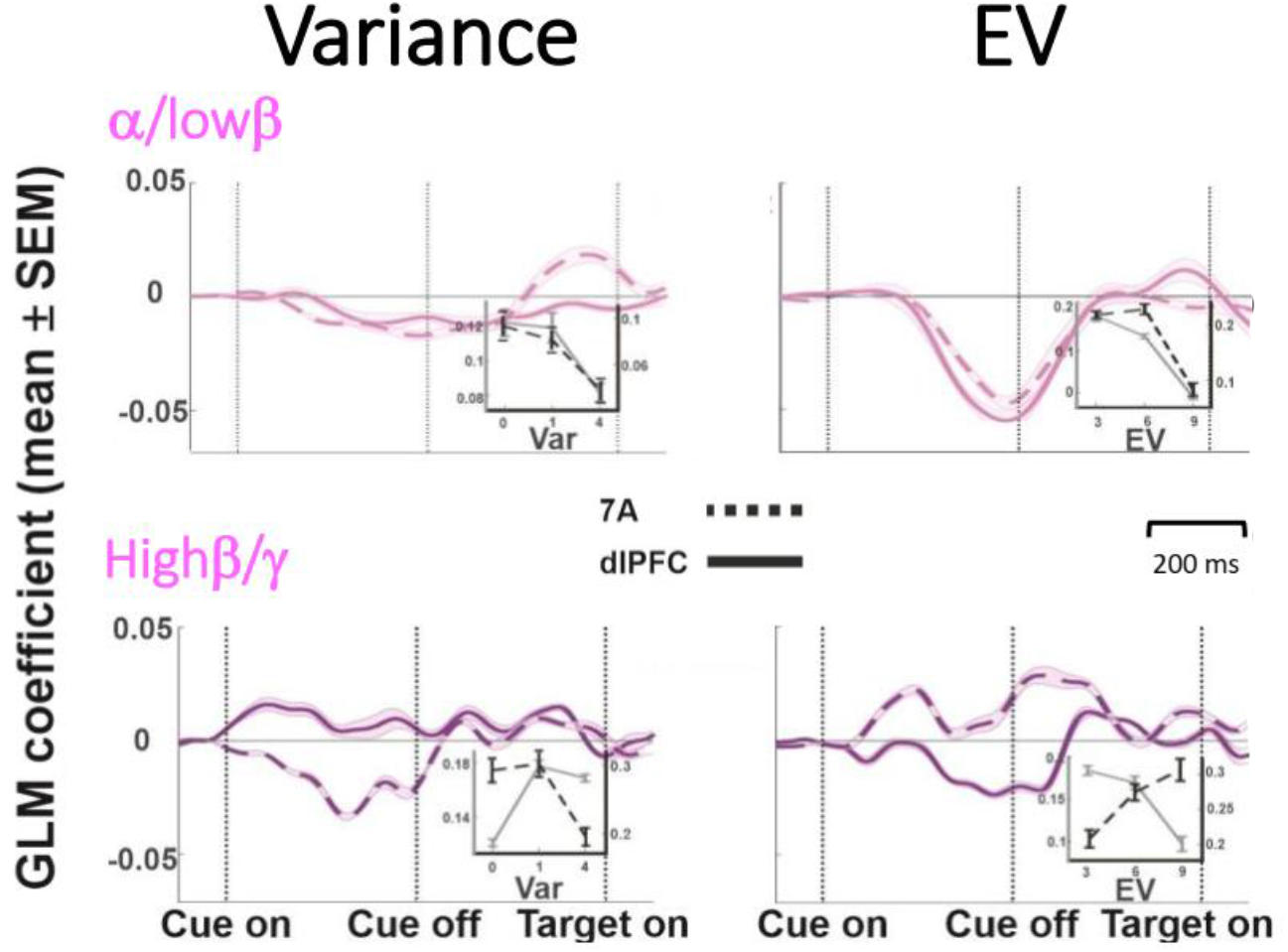
Variance and EV effects in the two ROIs. Each trace shows the average GLM coefficient in the ROIs shown in **Fig. 4** (mean and SEM across the frequencies included in the ROI). The insets show the raw LFP power in the time-frequency ROI as a function of variance or EV. The left gray axis and gray curve corresponds to dlPFC, and, the right black axis and dashed black curve corresponds to 7A. For these plots, z-transformed LFP power was averaged for each electrode, and the data show average and SEM across n = 48 electrodes as a function of the variable of interest (variance and EV).

Power in a α/low-β frequency band (8 - 18 Hz) is widely associated with task engagement and arousal in different tasks and brain areas in humans and monkeys^32^. Consistent with this widely replicated result, activity in this band was suppressed by variance and EV in both 7A and the dlPFC (**Fig. 4**, pink ROIs). The strongest effects arose in the late cue and early delay periods (**Fig. 5**, top row) and were highly significant for both variables for each monkey (7A variance: monkey 1: p < 6×10^−6^ (Wilcoxon rank sum test relative to 0 across all pixels in the ROI); monkey 2: p < 2×10^−14^; EV: monkey 1: p < 2×10^−21^, monkey 2: p < 9×10^−13^; dlPFC variance: monkey 1: p < 6×10^−10^, monkey 2: p < 8×10^−8^; EV: monkey 1: p < 5×10^−28^, monkey 2: p < 2×10^−12^).

In contrast with the uniform suppression in the low frequency band, the effects in the high-β/low-γ differed for variance and EV and across the two areas (**Fig. 5**, bottom row and **Fig. 4**, purple ROI). In 7A, power in this band was suppressed by variance and enhanced by EV (**Fig 5,** dashed traces bottom left vs bottom right panels), while the dlPFC showed the opposite pattern – being enhanced by variance and suppressed by EV **(Fig. 5, solid traces,** bottom left vs right panels). Each effect was highly robust in each monkey (7A variance: monkey 1: p < 2×10^−13^, monkey 2: p < 4×10^−17^; 7A EV: monkey 1: p < 2×10^−16^, monkey 2: p < 4×10^−18^; dlPFC variance monkey 1: p < 3×10^−6^, monkey 2: p < 2×10^−22^; dlPFC EV: monkey 1: p < 4×10^−28^, monkey 2: p < 9.5×10^−4^). Thus, variance and EV reduced power in the α/low-β frequency range in both areas but had distinct area-specific effects in the high-β/low-β frequency range.

### Variance enhances parietal to frontal information transmission

Given theoretical predictions that uncertainty modulates the balance between top-down and bottom-up information transmission^4, 6^, we asked how variance and EV modulate functional interactions among the two areas. To this end, we calculated spike field coherence (SFC) using the method of Vinck et al. that is known to compensate for biases due to low spike counts and volume conduction ^33, 34^. The SFC measures the extent to which spikes arrive at a consistent phase of the LFP oscillations and provides an index of directional interactions. The SFC between spikes in area A and LFPs in area B measures the extent to which outputs from area A influence area B, while the SFC between spikes in area B and LFPs in area A measure the opposite interactions^33–35^ (see also *Methods*).

The most robust modulation we found was an asymmetric effect of variance on fronto-parietal SFC. Higher uncertainty was associated with enhanced SFC from 7A spikes to dlPFC LFPs, suggesting enhanced information transmission from 7A to the dlPFC (**Fig. 6A,** left). Conversely, higher variance was associated with reduced SFC in the opposite direction, suggesting reduced information transmission from dlPFC to 7A (**Fig. 6A,** right). These effects were consistent in both monkeys (**Fig. 6B**) and could not be explained by changes in LFP power, which had opposite signs in the two areas (**Fig. 4,5**) or by LFP-LFP coherence, which did not show consistent modulations with variance or EV (**Fig. S6**). The SFC modulations were unique to variance and to across-area communications, with only weak and inconsistent effects being produced by EV on SFC across areas (**Fig. S5A**) and by both variance and EV within areas (**Fig. S5B**).

**fig. 6.**
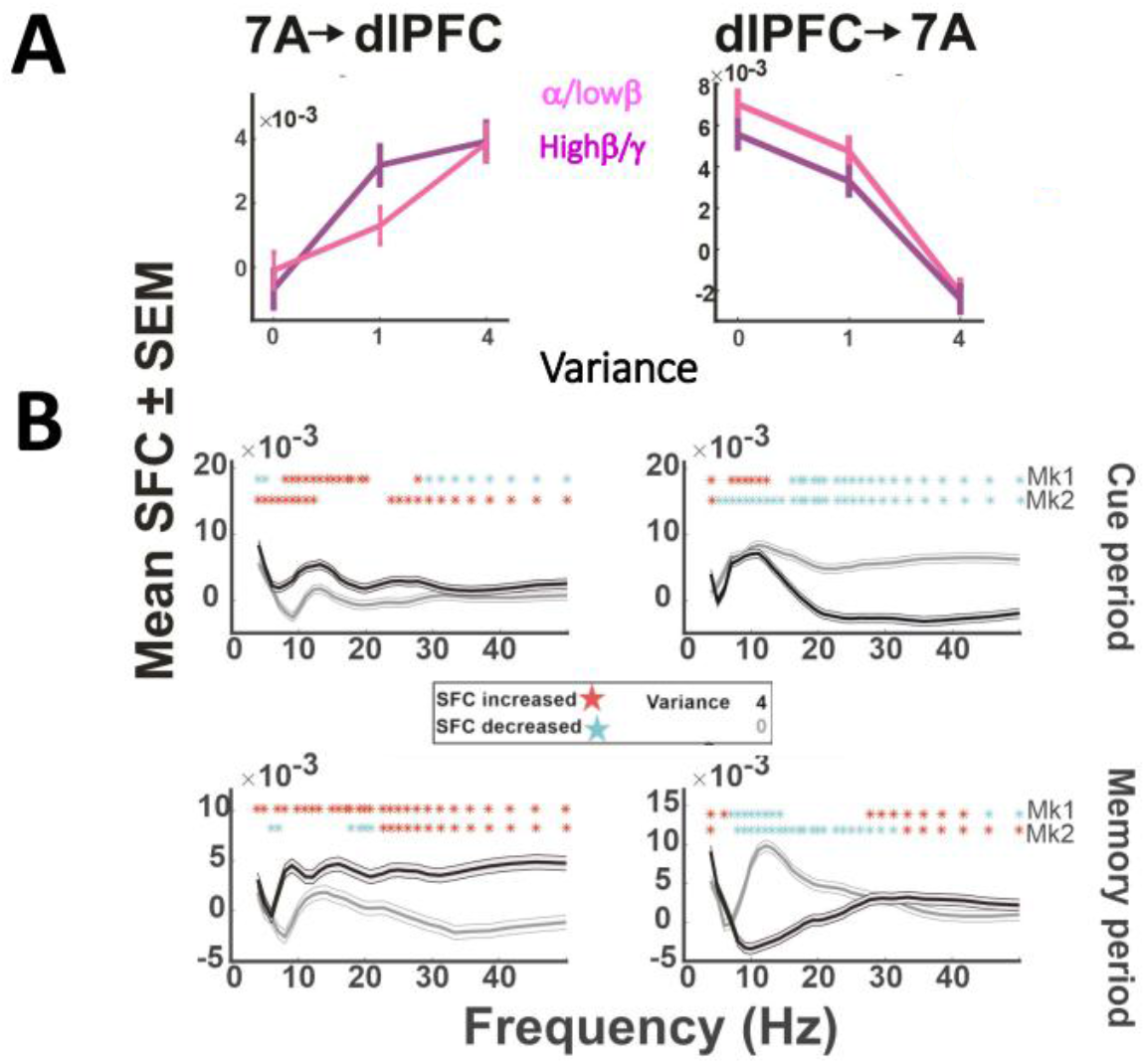
The effects of uncertainty on spike-field coherence (SFC) **(A) SFC as a function of variance** SFC as a function of variance in the time-frequency bins showing the strongest effects. Each point shows the mean and SEM of the SFC values across all neuron-LFP channel pairs and all task conditions with identical level of uncertainty. **(B) SFC as a function of frequency** The curves show the mean and SEM of the SFC values for different frequencies in the time bins that showed the most consistent effects (top row: 200-400 ms; bottom row: 400-600 ms (right) and 600-800 ms (left) after cue on). The stars in each panel show the frequencies where SFC was significantly modulated for each monkey (Kruskal-Wallis test, p<0.05 after correcting for multiple comparisons across frequencies). Blue stars indicate decrease and red stars indicate increase of SFC with variance. The SFC values for the peak effects in each panel are as follows (all units of 10^−3^). Cue period, mean±SEM SFC for 7A→dlPFC in the α/low-β frequency band: var_0_: −0.09±0.054, var_4_: 3.8±0.5; Kruskal- Wallis test, p = 5.2×10^−8^. For dlPFC→7A, in the high-β/low-β frequency band: var_0_: 5.5±0.6, var_4_: 2.4±0.6 (p = 1.8×10^−21^). Delay period, 7A→dlPFC in the high-β/low-β frequency band: var_0_: −0.7±0.6, var_4_: 9.3±0.6 (p = 3.5×10^−7^). dlPFC→7A, in the α/low-β frequency band: var_0_: 7.0±0.6, var_4_: −2.1±0.6 (p = 6.3×10^−16^).

The SFC modulations by variance extended to all frequency bands and differed across the task epochs (**Fig. 6B**). In the α/low-β frequency band, the earliest modulation was an increase in parietal to frontal SFC followed by a decrease in the frontal to parietal direction (**Fig. 6B**, upper left and lower right panels; all p < 10^−7^ in each monkey, Krusal-Wallis test; see figure legend for detailed statistics). In the high-β/low-γ frequency band this sequence was reversed, with the earliest modulation being reduction in frontal to parietal SFC followed by increased parietal to frontal SFC (**Fig. 6B**, upper right panel and lower left panel; all p < 10^−7^ in each monkey). Thus, uncertainty sets off an intricate temporal sequence of increases and decreases in fronto-parietal functional connectivity.

## Discussion

We show that reward uncertainty affects multiple aspects of microscopic and mesoscopic activity in monkey areas 7A and the dlPFC. Single-neuron responses, LFP oscillations and spike-field coherence were sensitive to the uncertainty of expected rewards, independently of the value of the rewards and the areas’ visuo-spatial selectivity.

One prominent effect of uncertainty was a reduction in α/low-β LFP power that was highly consistent in 7A and the dlPFC. Reductions in α/low-β LFP power have been widely reported across different tasks and brain structures and linked with enhanced task engagement, reduced inhibition, and desynchronized neural activity^36^. While some previously reported effects were spatially specific (correlated with processing of contralateral visual cues), a recent study found non-spatial reductions in α-band power at higher EV in monkey primary motor cortex^37^ consistent with the reductions we find (see also ^38^). Our findings thus show that reward-related effects on α/low-β LFP power extend to the fronto-parietal network and may mediate arousal produced by both EV and uncertainty.

In contrast with the homogeneous reduction in low-frequency power, uncertainty and EV produced much more heterogeneous modulations in the higher frequency range, which differed markedly in sign (enhancement or suppression) for each area and for variance versus EV. This result is consistent with the fact that β-band power has been variably reported to increase and decrease with attention across tasks and cortical areas^36^. Based on the prevailing view that γ-band oscillations primarily index feedforward sensory processing^30, 39^ our findings suggest that feedforward processing is differentially affected by uncertainty versus EV.

Another clear distinction we find is that variance and EV had separate representations in spiking activity. Previous studies have shown that reward variance is encoded independently of value by individual neurons in the orbitofrontal cortex^19^ and subcortical structures^20, 21^. Our findings show that this segregation extends to the fronto-parietal network and to circuit-based measures including noise correlations and population decoding capacity.

Importantly, rather than producing overall increases or decreases in firing, uncertainty and EV had opponent-coding representations, enhancing or suppressing responses in distinct classes of cells. While an opponent-coding representation has been previously reported for EV in the dlPFC^40, 41^ here we show that it extends to the parietal cortex and to uncertainty. In addition, our result that neurons with similar polarity have higher noise correlations gives strong support to the view that polarity defines distinct neural circuits with different behavioral functions. In the case of EV, neurons with positive and negative scaling may be associated with, respectively, approach and avoidance behaviors (go/no-go tendencies) that are mediated by distinct basal ganglia pathways^42^. By analogy, the opponent-coding of uncertainty suggests that the brain has distinct circuits for controlling behaviors in tasks with high and low uncertainty. One idea has been that the brain uses uncertainty to arbitrate between controllers of different complexity – relying on a simpler striatal controller when familiar, habitual strategies are sufficient but engaging the prefrontal cortex in uncertain conditions^43^. An interesting hypothesis, therefore, is that uncertainty sensitive cells with positive- and negative-scaling may be part of brain-wide circuits mediating distinct forms of control.

A central result we report is that uncertainty had powerful effects on fronto-parietal connectivity. Higher uncertainty was associated with reduced SFC from the frontal to the parietal cortex but *enhanced* SFC from the parietal to the frontal lobe. These results are consistent with a recent report that, although fronto-parietal areas have similar single-neuron activity, the direction of their functional interactions can be strongly dependent on context^26^. That study found that, in monkeys performing a familiar categorization task, information about task context and rules was predominantly transmitted in a top-down direction, from dlPFC to 7A. Our findings confirm this result by showing that, when monkeys have low uncertainty, frontal-to-parietal SFC is higher than parietal-to-frontal SFC (i.e., 0 variance, **Fig. 6A**). We extend this result by showing that this balance parametrically depends on uncertainty, consistent with computational theories of optimal inference.

Our finding that uncertainty reduces SFC in the top-down direction does not imply that the dlPFC goes “offline” in uncertain conditions. Indeed, the modulations of frontal-to-parietal and parietal-to-frontal SFC had overlapping time-courses. Moreover, some of the strongest effects of uncertainty – i.e., on high-β/low-γ frequency LFP power (**Fig. 4**) and SFC (**Fig. 6B**) - emerged first in the dlPFC and only later in 7A. It is thus possible that, consistent with current models of executive function, uncertainty is detected by frontal cortical areas including the dlPFC and the anterior cingulate cortex and these areas provide the initial drive which, perhaps by triggering release of neuromodulators, ultimately leads to increases in sensory gains and enhancements in parietal to frontal transmission^2, 44^.

Our findings also support the idea that, while the parietal cortex is not a strictly sensory area, it plays an important role in resolving uncertainty. Early support for this view comes from the reinforcement learning literature showing that rats have increases in associability (learning rates) for uncertain sensory cues and these increases are reduced by lesions of the parietal cortex^45^. Subsequent single-neuron recordings in monkeys provide additional support for this view both indirectly, by revealing enhanced parietal responses to novel stimuli and salient distractors^16, 46^ and directly, by showing that, in multi-step decision tasks, parietal cells assign credit and learning specifically at junctures that resolve uncertainty^11, 47, 48^. Thus, an important direction for future research is to refine our understanding of the intricate mechanisms that allow the brain to allocate resources to task junctures that are not only associated with consequential outcomes but have high uncertainty and benefit from cognitive strategies that resolve the uncertainty^5, 7, 18^.

## Methods

### General methods

Data were collected from two adult male rhesus monkeys (Macaca mulatta; 9-12kg) using standard behavioral and neurophysiological techniques as described previously (Oristaglio et al. 2006). All methods were approved by the Animal Care and Use Committees of Columbia University and New York State Psychiatric Institute as complying with the guidelines within the Public Health Service Guide for the Care and Use of Laboratory Animals. Visual stimuli were presented on a MS3400V XGA high definition monitor (CTX International, INC., City of Industry, CA; 62.5by 46.5 cm viewing area). Eye position was recorded using an eye tracking system (Arrington Research, Scottsdale, AZ). Licking was recorded using an in-house device that detected interruptions in a laser beam produced by extensions of the monkeys’ tongue to obtain water rewards.

### Task

A trial started with the presentation of two textured square placeholders (1 ° width) located along the horizontal meridian at 8° eccentricity to the right and left of a central fixation point (white square, 0.2° diameter). After a 300-500 ms period of central fixation (when the monkeys maintained gaze within a 1.5-2° square window centered on the fixation point) one of the placeholders was replaced by a randomly selected reward cue (a vertical rectangle measuring 1.2 × 5° with 11 gray bars indicating the reward scale, and one or two gradations highlighted in yellow, indicating the trial’s rewards). The cue remained visible for 400 ms and was followed by a 400ms delay period, after which the fixation point disappeared and one of the placeholders simultaneously increased in luminance, indicating the saccade target. The target location was randomized independently of the reward cue. If the monkey made a saccade to the target with a reaction time (RT) of 100 ms – 700 ms and maintained fixation within a 2.0-3.5° window of it for 377 ms, he received a reward with the magnitude and probability that had been indicated by the cue.

### Neural recordings

After completing behavioral training, each monkey was implanted with two 48-electrode Utah arrays (electrode length 1.5 mm) arranged in rectangular grids (1 mm spacing; monkey 1, 7×7 mm, monkey 2, 5×10 mm) and positioned in the pre-arcuate portion of the dlPFC and the posterior portion of area 7A (**Figure S1**). Data were recorded using the Cereplex System (Blackrock, Salt Lake City, Utah) over 24 sessions spanning 4 months after array implantation in monkey 1, and 11 sessions spanning 2 months after implantation in monkey 2.

### Data analysis

We analyzed a total of 12,029 trials that (1) were correctly completed and (2) had RT within 2 standard deviations relative to the mean of each monkey’s full data set (monkey 1: n = 8,082 analyzed trials, monkey 2: n = 3,947. Data were analyzed with MatLab (MathWorks, Natick, MA; version R2016-b) and other specialized software as noted below.

### Licking

The lickometer signal was digitized at 1 kHz to produce a trial by trial record of licking with 1 ms resolution. The probability of licking was measured in a 400 ms time window centered on the time of each monkey’s average peak licking response (monkey 1: 400-800 ms after cue onset; monkey 2: 800-1,100 ms after cue onset). Licking probabilities in individual trials were normalized by subtracting the session’s mean, pooled across sessions and subjected to a generalized linear model (GLM) analysis with EV and variance, including cue position and the EV×variance interaction as nuisance regressors (using a binomial distribution and logit link function and implemented in the fitglm function in the MATLAB statistics toolbox). Models that included a parametric uncertainty regressor outperformed those that included only a binary indicator of probabilistic versus deterministic cues and are presented throughout the paper.

### Single neurons spike analysis

Raw spikes were sorted offline using WaveSorter and produced a total of 1,175 neurons in dlPFC (749 in monkey 1) and 971 neurons in 7A (755 in monkey 1). We focused the analysis on the subset of units that were well isolated, had at least 5 trials in each condition, and fired at least 5 spikes on average within the time interval from 500 ms before to 1,000 ms after cue onset, comprising 530 neurons in dlPFC (432 in monkey 1) and 522 neurons in 7A (481 in monkey 1).

To measure neuronal selectivity, we fit each neuron’s trial-by-trial spike count in the interval 0 – 800 ms after cue onset using a GLM with factors EV, variance, EV× variance, and Cue location, using a normal distribution and identity link function. To estimate changes in firing rate variability, we computed the Fano factor – the ratio of across-trial variability to the mean firing rate. Although the Fano factor was lower during the cue/delay relative to the pre-cue epochs, we found no consistent changes as a function of variance or EV in either area.

### SVM classification

We smoothed the raw spike train using a Gaussian kernel of 50 ms standard deviation and measured the average smoothed firing rate in each trial in the interval 0 – 800 ms after cue onset. We then evaluated decoding accuracy for each pairwise classification (e.g., EV3 vs EV6, variance 1 vs variance 4, etc) using the data pooled across all the neurons in an array. To construct the pooled dataset, we randomly selected *m* trials from each neuron and every condition, where *m* was equal to the minimum number of trials across all neurons and all conditions. We used a 5-fold cross-validation procedure with 200 repetitions to compute decoding accuracy in the original data set and repeated the procedure after randomly shuffling trial labels to compute the baseline accuracy expected purely by chance.

### LFP pre-processing

The raw local field potentials (LFP) from each electrode and trial were measured from 1,200 ms before to 2,000 ms after cue onset, notch filtered at 60Hz, low pass filtered at 100Hz and subjected to a linear trend removal. The traces from each session were then pooled and subjected to a two-step cleaning procedure to remove outliers in, respectively, the frequency and time domains. For the first step that removed outliers in the frequency domain, we calculated the power spectrum of each LFP trace in the range of 0 - 90 Hz (using a multi-taper method with 4 tapers) and characterized each trial with a 5-dimensional vector containing the sum of the logarithm of the power spectrum in 5 frequency bands (0.5-4 Hz, 4-8 Hz, 8-12 Hz, 12-30 Hz and 30-90 Hz). We then reduced the dimensionality of each session’s data set to 2 principal components using principal component analysis and clustered this 2- dimensional data set using Gaussian Mixture Models (GMM; *fitgmdist* function in the MATLAB *statistics and machine learning toolbox*). This procedure produced, for each session, one or two “dense” clusters that contained most of the session’s data, and one or two “sparse” clusters containing the remaining trials, in which the LFP power in at least one frequency band was an outlier. We discarded the trials in the sparse clusters. In addition, we discarded trials that were identified as outliers within the dense clusters - i.e., for which the Mahalanobis distance to all other trials in the cluster was above the 90th percentile. The trials surviving the first step were subjected to a second step that removed outliers in the time domain. To this end, we computed the peak-to-peak amplitude of the broadband LFP trace in each trial, z transformed these values, and removed trials for which this measure was more than half a standard deviation away from the mean across all trials. This was a conservative cleaning procedure that removed all the trials with poor signal quality due to a variety of reasons (e.g., signal to noise ratio, artefact or saturation). Overall, 39.5% of trials (12.3% to 77.4% across sessions) were excluded after preprocessing.

### LFP power spectrum

For each trial that was accepted for analysis, we calculated the LFP power spectrum in 1 Hz frequency bands using Morlet wavelet transformation (*ft_freqanalysis* function of the FieldTrip toolbox^49^. The power in each band was then z-scored relative to all the trials and time points within the session, and normalized relative to the trial’s baseline using the following equation:

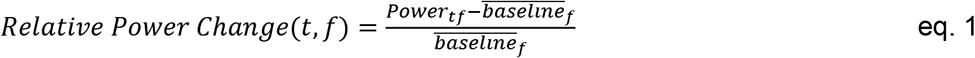

where *Power*_*tf*_ is the power at time *t* and frequency *f*, and 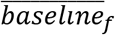 is the power in frequency *f* during the 300 ms time interval before cue onset on the same trial. By normalizing relative to the frequency-specific baseline, this procedure accounts both for trial by trial variability and 1/f power distribution^34^.

### GLM of LFP power spectrum

The *Relative Power Change* quantity from eq. 1 produced a time-frequency map of normalized LFP power for each trial and electrode. To determine how these maps varied as a function of uncertainty and EV, we pooled the trials across sessions and electrodes within each array, and fit this pooled dataset using a GLM with factors of [EV, variance, EV×variance, Cue location] assuming a normal distribution and identity link function. This produced a time-frequency map of coefficients measuring the effects of EV and variance, controlling for any visuo-spatial response and EV×variance interaction(**Fig. 3**). To identify regions of interest within the GLM coefficient maps, we divided the cue and delay periods into 200 ms epochs, and identified frequencies for which the coefficients for a variable were significantly different from 0 with the same sign in both monkeys (Kruskal-Wallis test with False Discovery Rate (FDR) correction).

### Field-field coherence

was measured using weighted phase lag index (WPLI)^35^. The WPLI uses imaginary part of the cross spectrum to remove the volume conduction effect. Within a session, for every task condition the phase locking index was calculated across trials and LFP channel pairs for every time and frequency. GLMs with EV, variance and EV×variance factors were then fitted to the coherence maps from different sessions, assuming normal distribution and identity link function.

### Spike-field coherence

We used the FieldTrip toolbox^49^ to calculate the power spectrum for the trial-by-trial LFP using multitaper analysis, and the *ft_spiketriggeredspectrum* function to measure the phase in frequencies of 4-47 Hz. We estimated SFC using the average Pairwise Phase Consistency index (PPC2, *ft_spiketriggeredspectrum_stat* function), which is known to minimize biases due to low spike counts and volume conduction^33, 35^. For every pair of neuron-LFP channel, in each task condition and frequency, PPC2 was calculated across all spikes that the cell fired in the corresponding task condition. PPC2 values of all cell-LFP pairs (excluding pairs in which the neurons did not emit any spikes) were submitted to non-parametric analyses to detect influences of EV and variance (n ranging between 136,368 and 164,880 across conditions). In the frequency plots, p values were corrected for comparison across frequencies using False Discovery Rate correction method.

## Acknowledgements

The work was supported by a generous gift from Synthes, Inc. and a McKnight Foundation Memory and Cognitive Disorder Award to JG.

## Data availability

The data recorded and analyzed in this study are available from the corresponding author upon reasonable request.

---

**Figure S1.**
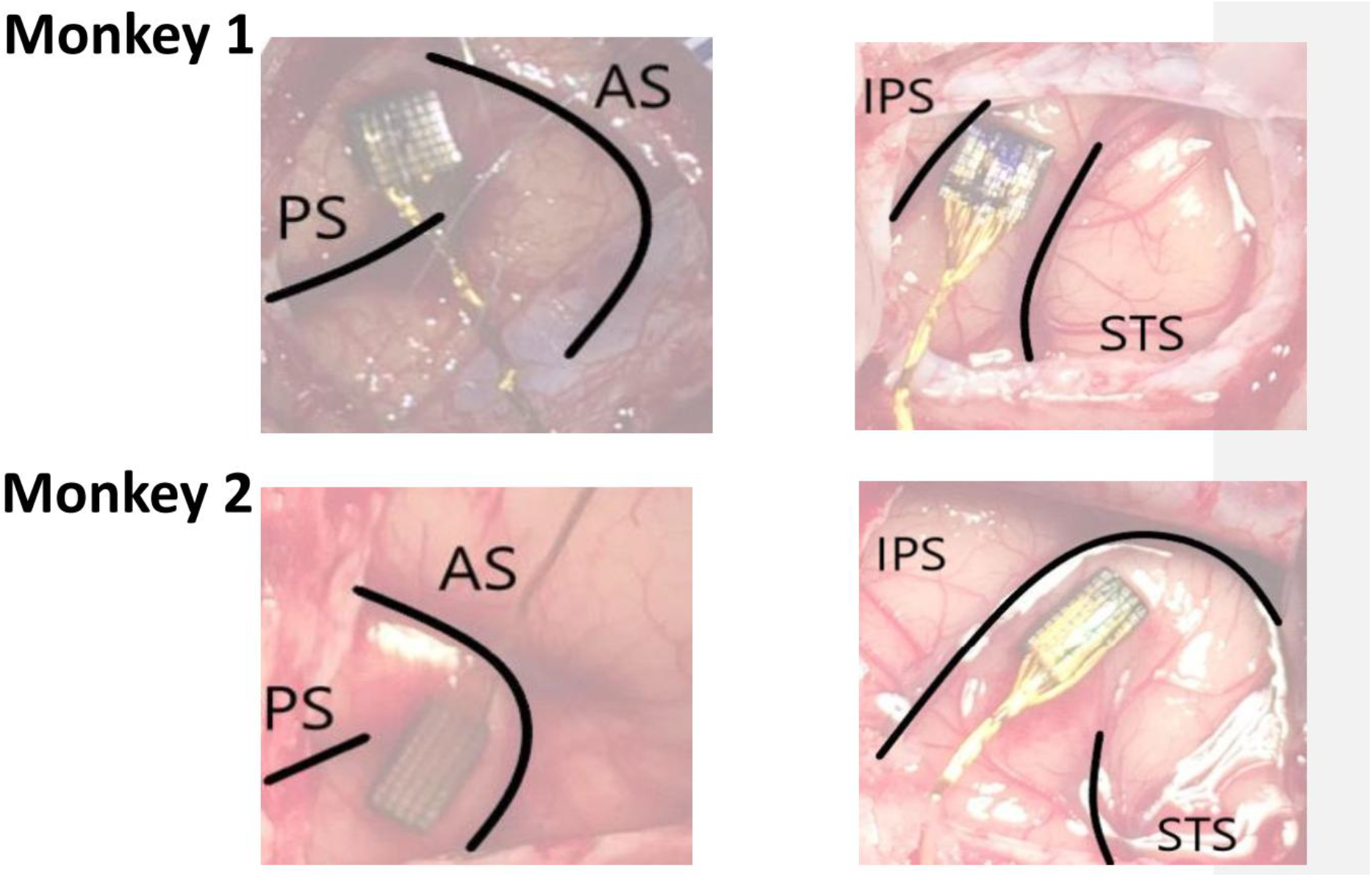
Recording sites. lntraoperative photographs showing array placements. **(A)** The dlPFC arrays were implanted between the arcuate sulcus (AS) and the principal sulcus (PS), slightly more dorsal in monkey 1 relative to monkey 2 because of vascular anatomy. **(B)** The 7A arrays were implanted between the intraparietal sulcus (IPS) and superior temporal sulcus (STS), in the posterior portion of this area that has been targeted in recent studies.

**Fig. S2.**
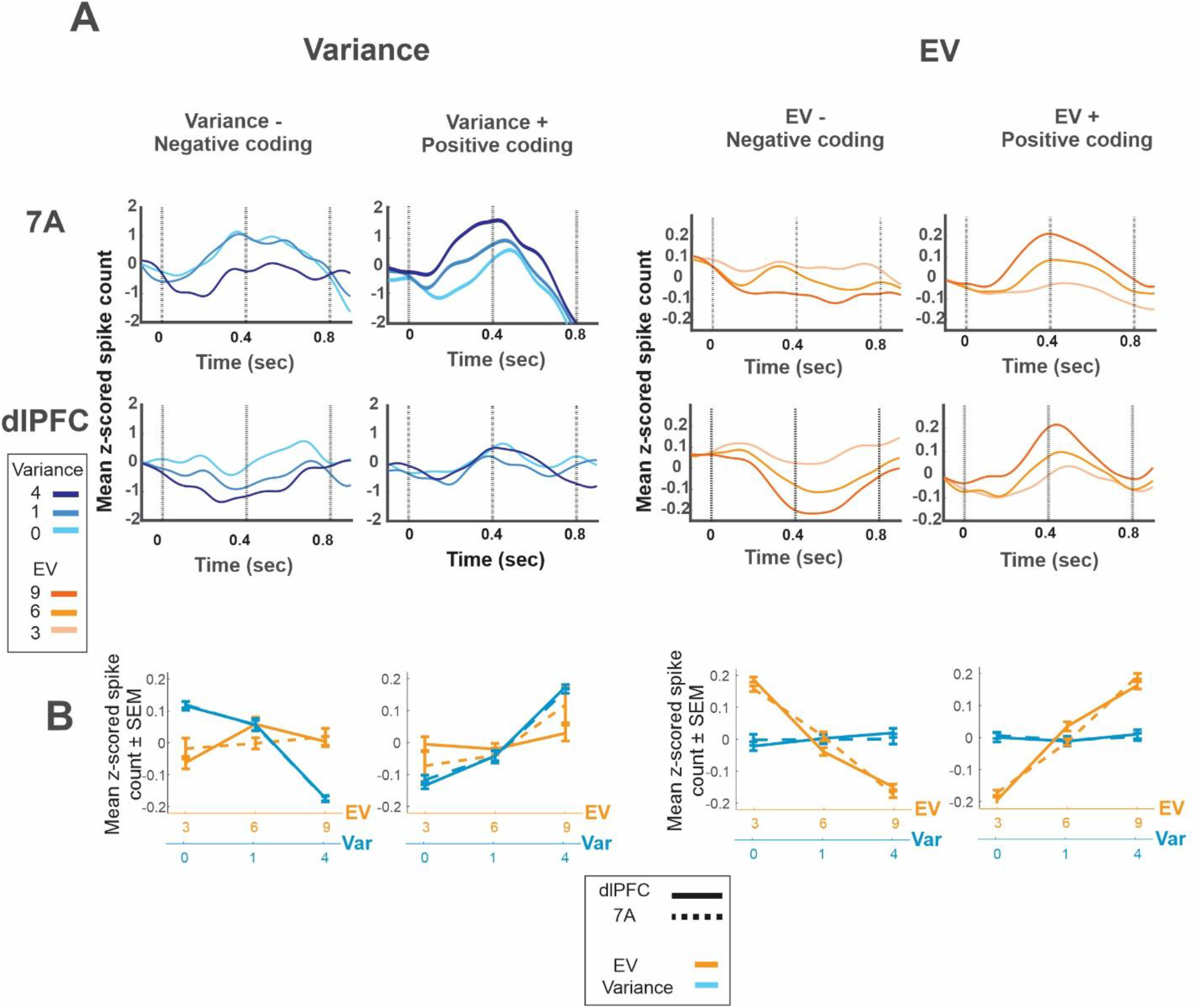
Positive and negative encoding of variance and EV. **(A) peri-stimulus time histograms** for the subsets of cells with significant positive (columns 2 and 4) or negative coefficients (columns 1 and 3) for variance and EV (including the few cells with both main effects). The traces show average z-scored spike counts across cells, aligned on cue onset, smoothed with a Gaussian kernel with 50 ms standard deviation, and sorted by variance (two left columns) and EV (two right columns). **(B) Spike counts** of the sensitive cells shown in A as a function of variance (blue) and EV (orange), for dlPFC (solid) and 7A (dashed). Each point shows the mean of the spike count in the interval of 0-800 ms after cue onset, z-scored within each cell and averaged across cells. Error bars show SEM across cells.

**Fig. S3.**
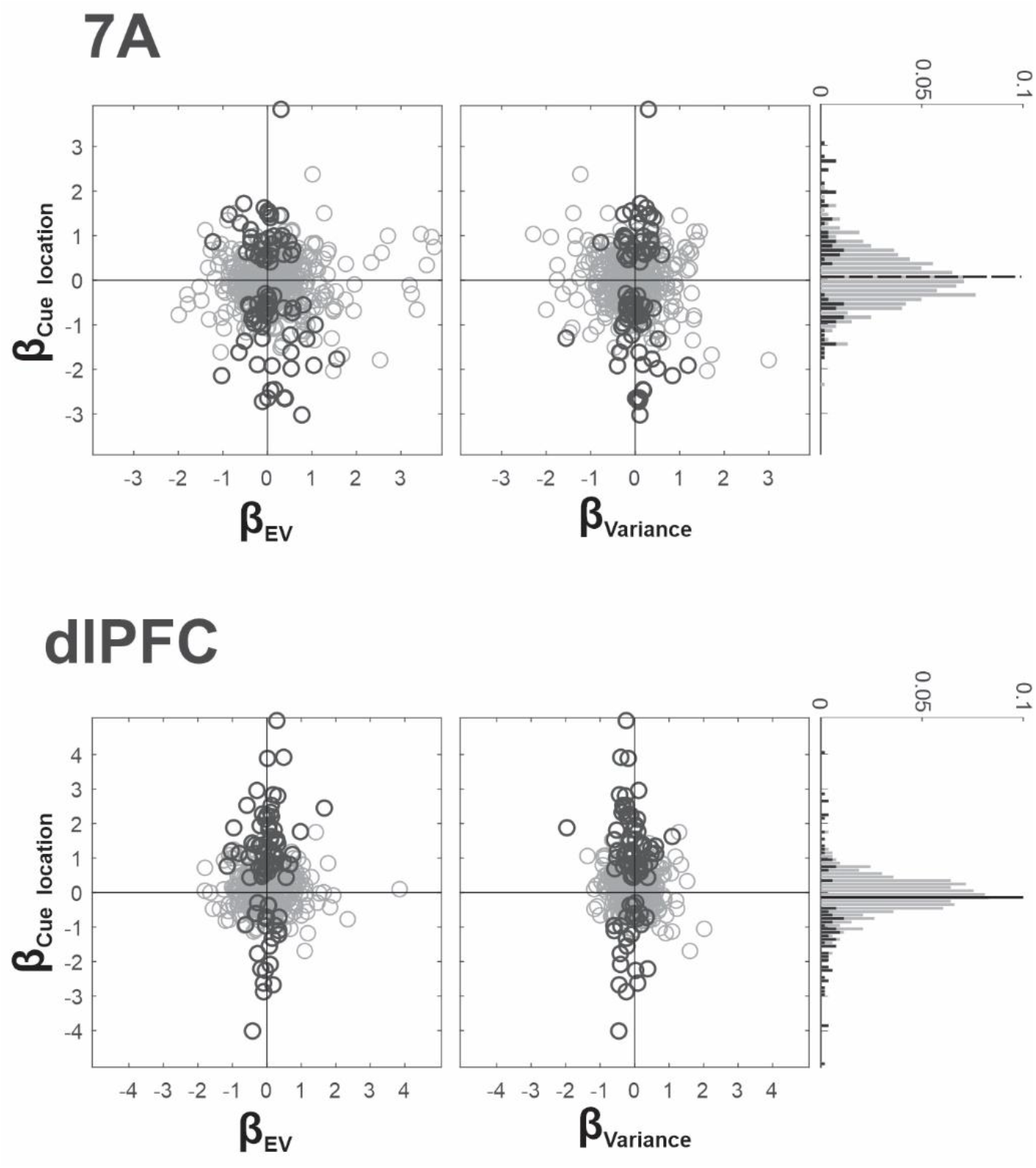
Encoding of cue location in single-neuron spiking activity comparing GLM coefficients for the cue location (β_Cue location_, ordinate) with coefficients of EV (β_EV_, left panel, abscissa) and variance (β_Variance_, right panel, abscissa) in individual neurons of the 7A and dlPFC. Every point is one neuron and dark color indicates cells with significant coefficients for cue location. Marginal distribution shows the significant coefficients in black, with the median indicated by a vertical line (solid line if the median is significantly different from 0, p < 0.05, Wilcoxon signed-rank test, and white otherwise). Across the population of all cells there was a bias towards contralateral encoding in dlPFC (mean ± SEM coefficients 0.134 ± 0.035; p = 25×10^−5^) but not in 7A (mean ± SEM coefficients −0.059 ± 0.032; p = 0.42). Cue location encoding was not correlated with either variance or EV encoding (Spearman correlation, all p>0.06).

**Fig. S4.**
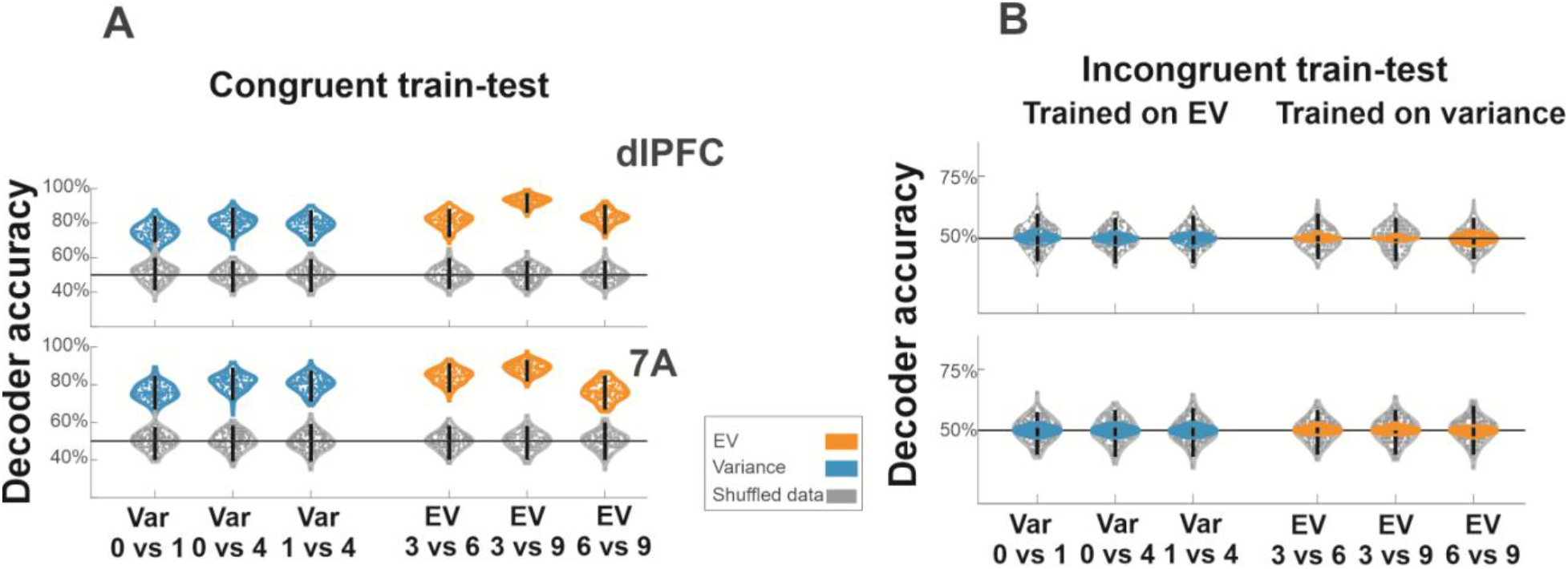
Decoding accuracy for congruent and incongruent train-testing regimes. **(A) Congruent regime.** Each violin plot shows the test accuracy scores for binary classification of EV (orange) or variance levels (blue) for classifiers trained on the same variable. Each violin plot shows the accuracy over 200 bootstrap iterations from the original dataset (colored) and the dataset with randomized labels (gray). The whiskers in each distribution show 95% confidence interval around the mean. Accuracy is above change if the CI do not include zero and do not overlap with the randomized distributions. **(B) Incongruent regime.** Same as in A, but for classifiers trained and tested on the other variable.

**Fig. S5.**
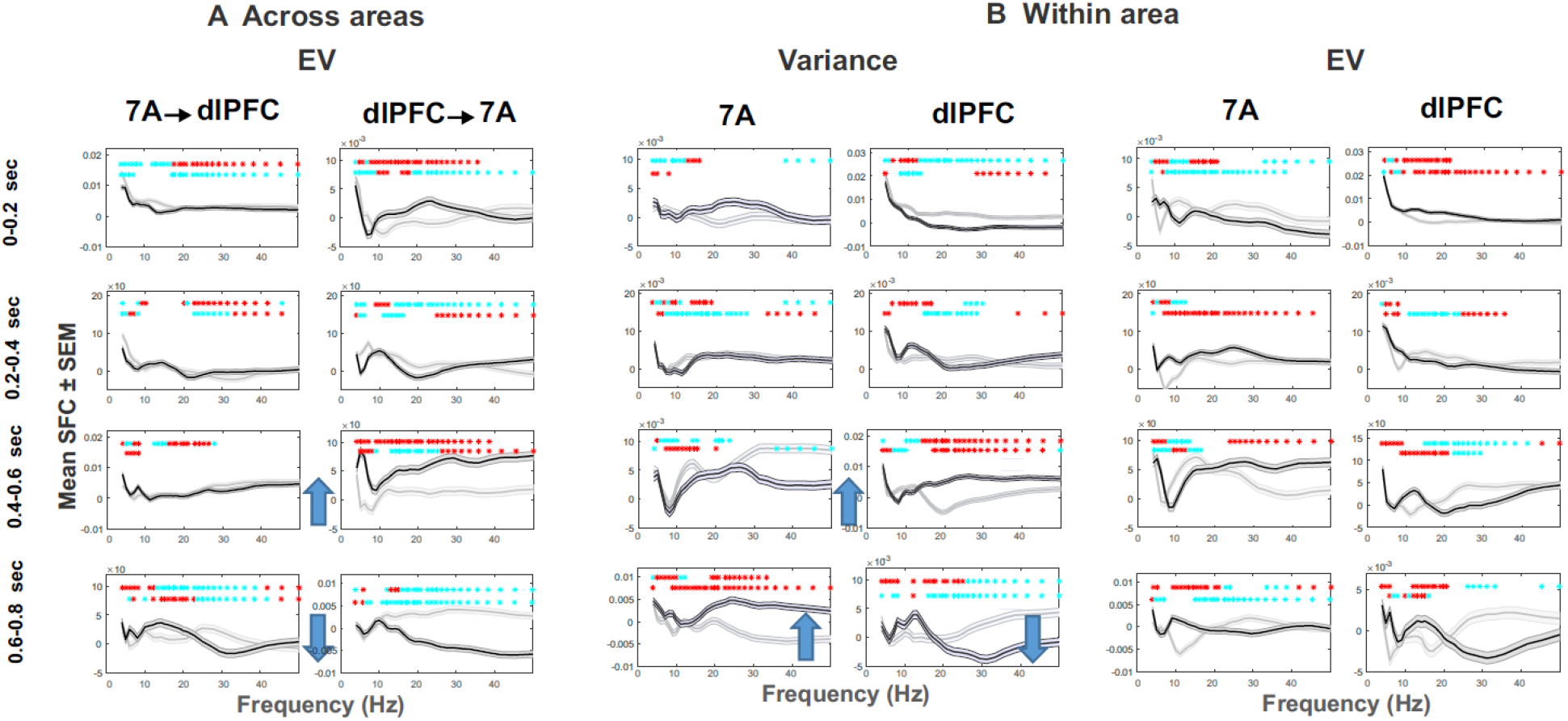
SFC is not consistently modulated by EV across areas. **(A) or by variance or EV within area (B)** From top to bottom, rows show successive 200 ms intervals aligned on cue onset. Blue arrows show intervals in which both monkeys showed significant modulations of the same sign. In **A**, this included an increase followed by decrease in dlPFC to 7A SFC in response to EV in, respectively, the early and late memory periods. In **B**, it included an increase followed by decrease in within-dlPFC SFC in response to variance in, respectively, the early and late memory periods. All other conventions as in **Fig 6B**.

**Fig. S6.**
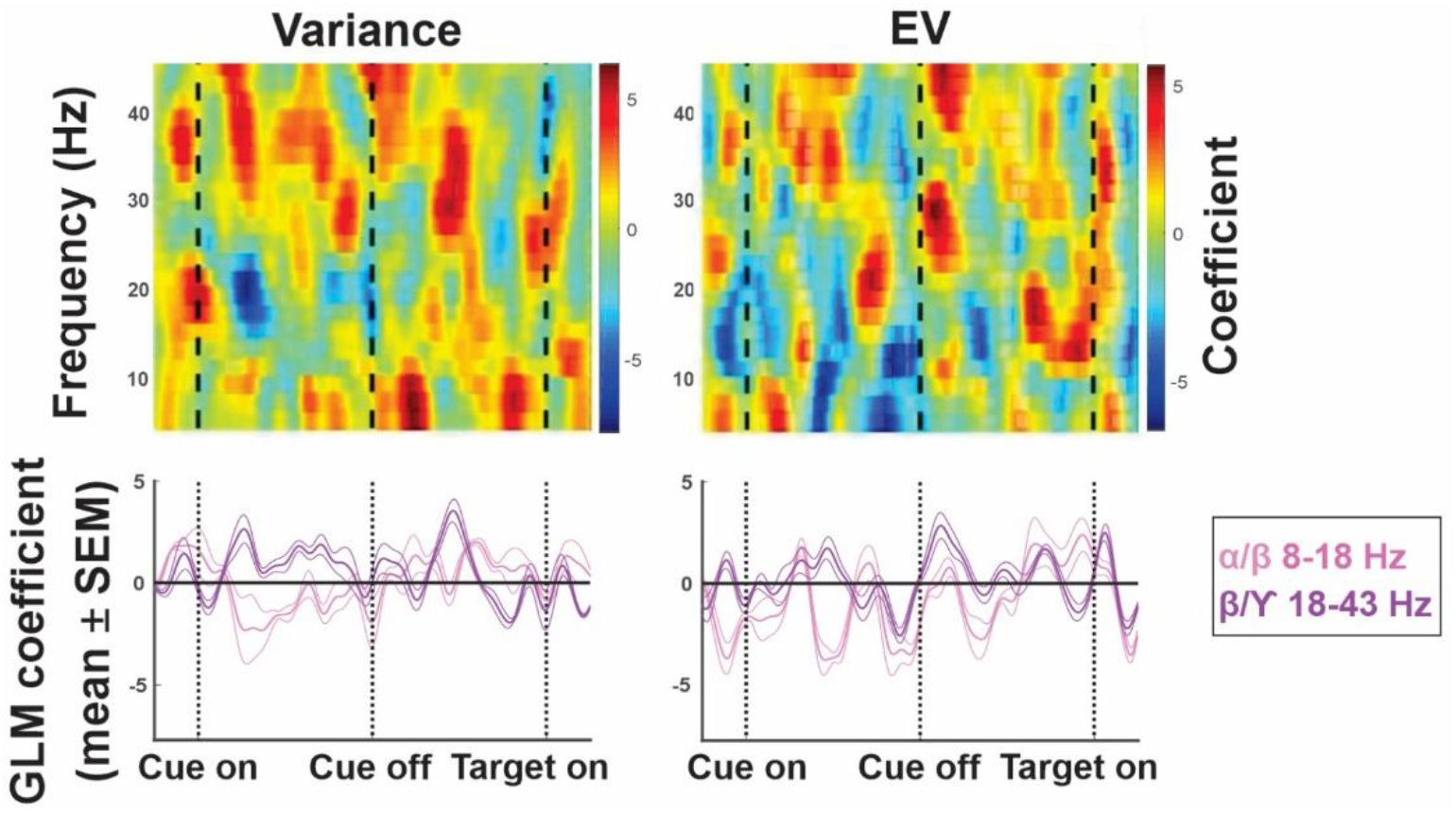
LFP-LFP coherence. No consistent modulation with variance or EV. The degree to which two areas synchronize their oscillations is a sensitive index of neural connectivity. We therefore tested whether, and how, the coherence of oscillations between frontal and parietal LFPs was affected by variance and EV. In every session, for different time and frequencies (1 ms time bins and 2 Hz frequency bins), Weighted Phase Lag Index (WPLI; debiased WPLI option in the ft_connectivityanalysis() function of the FieldTrip toolbox) of LFP was calculated across trials and all electrode pairs between dlPFC and 7A. GLM with factors of EV, variance and their interaction was then fitted to the coherence maps from different sessions, assuming normal distribution and identity link function. There was no consistent strong modulation of coherence with variance or EV.

## References

1. Shenhav, A., Botvinick, M. & Cohen, J. The expected value of control: an integrative theory of anterior cingulate cortex function. Neuron 79, 217–240 (2013).

2. Shenhav A et al. Toward a Rational and Mechanistic Account of Mental Effort. Annu Rev Neurosci. 40, 99–124 (2017).

3. Grossberg, S. Adaptive pattern classification and universal recoding, II: Feedback, expectation, olfaction, and illusions. Biological Cybernetics, 23, 187–202 (1976).

4. Yu, A.J. & Dayan, P. Uncertainty, neuromodulation, and attention. Neuron 46, 681–692 (2005).

5. Gottlieb, J. & Oudeyer, P.Y. Toward a neuroscience of active sampling and curiosity. Nat Rev Neurosci. 19, 758–770 (2018).

6. Pezzulo, G., Rigoli, F. & Friston, K.J. Hierarchical Active Inference: A Theory of Motivated Control. Trends Cogn Sci 22, 294–306 (2018).

7. Fan, J. An information theory account of cognitive control. Front Hum Neurosci. 8(2014).

8. Kidd, C. & Hayden, B.Y. The psychology and neuroscience of curiosity. Neuron 88(2015).

9. Baranes, A.F., Oudeyer, P.Y. & Gottlieb, J. Eye movements encode epistemic curiosity in human observers. Vis Res in press(2015).

10. Daddaoua, N., Lopes, M. & Gottlieb, J. Intrinsically motivated oculomotor exploration guided by uncertainty reduction and conditioned reinforcement in non-human primates. Sci Rep 6(2016).

11. Horan, M., Daddaoua, N. & Gottlieb, J. Parietal neurons encode information sampling based on decision uncertainty. Nat Neurosci 22, 1327–1335 (2019).

12. Bach, D.R. & Dolan, R.J. Knowing how much you don’t know: a neural organization of uncertainty estimates. Nat Rev Neurosci 13, 572–586 (2012).

13. Platt, M.L. & Huettel, S.A. Risky business: the neuroeconomics of decision making under uncertainty. Nat Neurosci 11, 398–403 (2008).

14. Grabenhorst, F., Tsutsui, K.I., Kobayashi, S. & Schultz, W. Primate prefrontal neurons signal economic risk derived from the statistics of recent reward experience. Elife 8(2019).

15. Ebitz, R.B., Albarran, E. & Moore, T. Exploration disrupts choice-predictive signals and alters dynamics in prefrontal cortex. Neuron 97, 450–461 (2018).

16. Foley, N.C., Jangraw, D.C., Peck, C. & Gottlieb, J. Novelty enhances visual salience independently of reward in the parietal lobe. J neurosci 34, 7947–7957 (2014).

17. Hayden, B.Y., Heilbronner, S.R., Pearson, J.M. & Platt, M.L. Surprise signals in anterior cingulate cortex: neuronal encoding of unsigned reward prediction errors driving adjustment in behavior. Journal of Neuroscience 31, 4178–4187 (2011).

18. Gottlieb, J., Hayhoe, M., Hikosaka, O. & Rangel, A. Attention, reward and information seeking. Journal of Neuroscience 34, 15497–154504 (2014).

19. O’Neill, M. & Schultz, W. Coding of reward risk by orbitofrontal neurons is mostly distinct from coding of reward value. Neuron 68, 789–800 (2010).

20. Monosov, I.E. & Hikosaka, O. Selective and graded coding of reward uncertainty by neurons in the primate anterodorsal septal region. Nat Neurosci 16, 756–762 (2013).

21. Monosov, I.E., Leopold, D.A. & Hikosaka, O. Neurons in the Primate Medial Basal Forebrain Signal Combined Information about Reward Uncertainty, Value, and Punishment Anticipation. Journal of Neuroscience 35, 7443–7459 (2015).

22. Leong, Y., Radulescu, A., Daniel, R., DeWoskin, V. & Niv, Y. Dynamic Interaction between Reinforcement Learning and Attention in Multidimensional Environments. Neuron 93, 451–463 (2017).

23. Padmala, S. & Pessoa, L. Reward reduces conflict by enhancing attentional control and biasing visual cortical processing. J Cogn Neurosci 23, 3419–3432 (2011).

24. Kennerley, S.W. & Wallis, J.D. Reward-dependent modulation of working memory in lateral prefrontal cortex. J Neurosci 29, 3259–3270 (2009).

25. Mansouri, F.A., Egner, T. & Buckley, M.J. Monitoring Demands for Executive Control: Shared Functions between Human and Nonhuman Primates. Trends Neurosci 40, 15–27 (2017).

26. Crowe, D.A. et al. Prefrontal neurons transmit signals to parietal neurons that reflect executive control of cognition. Nat Neurosci 16, 1484–1491 (2013).

27. Katsuki, F. & Constantinidis, C. Early involvement of prefrontal cortex in visual bottom-up attention. Nat Neurosci 15, 1160–1166 (2012).

28. Cohen, M.R. & Kohn, A. Measuring and interpreting neuronal correlations. Nat Neurosci 14, 811–819 (2011).

29. Haegens, S. & Zion Golumbic, E. Rhythmic facilitation of sensory processing: A critical review. Neurosci Biobehav Rev 86, 150–165 (2018).

30. Bastos, A.M. et al. Visual areas exert feedforward and feedback influences through distinct frequency channels. Neuron 85, 390–401 (2015).

31. Bastos, A.M., Vezoli, J. & Fries, P. Communication through coherence with inter-areal delays. Curr Opin Neurobiol 31, 173–180 (2015).

32. Van Diepen, R.M., Foxe, J.J. & Mazaheri, A. The functional role of alpha-band activity in attentional processing: the current zeitgeist and future outlook. Curr Opin Psychol 29, 229–238 (2019).

33. Vinck, M., Battaglia, F.P., Womelsdorf, T. & Pennartz, C. Improved measures of phase-coupling between spikes and the local field potential. J. Comput. Neurosci. 33, 53–75 (2012).

34. Cohen, M.X. Analyzing neural time series data: theory and practice. (MIT Press, 2014).

35. Martin Vinck, M., Oostenveld, R., van Wingerden, M., Battaglia, F. & Pennartz, C.M.A. An improved index of phase-synchronization for electrophysiological data in the presence of volume-conduction, noise and sample-size bias NeuroImage 55, 1548–1565 (2011).

36. Thiele, A. & Bellgrove, M.A. Neuromodulation of Attention. Neuron 97, 769–785 (2018).

37. An, J., Yadav, T., Hessburg, J.P. & Francis, J.T. Reward Expectation Modulates Local Field Potentials, Spiking Activity and Spike-Field Coherence in the Primary Motor Cortex. eNeuro 6(2019).

38. Lemaire, N. et al. Effects of dopamine depletion on LFP oscillations in striatum are task-and learning-dependent and selectively reversed by L-DOPA. Proc Natl Acad Sci U S A 109, 18126–18131 (2012).

39. Buffalo, E.A., Fries, P., Landman, R., Buschman, T.J. & Desimone, R. Laminar differences in gamma and alpha coherence in the ventral stream. Proc Natl Acad Sci U S A 108, 11262–11267 (2011).

40. Kennerley, S.W., Behrens, T.E. & Wallis, J.D. Double dissociation of value computations in orbitofrontal and anterior cingulate neurons. Nat Neurosci. 14, 1581–1589 (2012).

41. Wallis, J.D. & Kennerley, S.W. Heterogeneous reward signals in prefrontal cortex. Current opinion in neurobiology 20, 191–198 (2010).

42. Calabresi, P., Picconi, B., Tozzi, A., Ghiglieri, V. & Di Filippo, M. Direct and indirect pathways of basal ganglia: a critical reappraisal. Nat Neurosci 17, 1022–1030 (2014).

43. Daw, N.D., Niv, Y. & Dayan, P. Uncertainty-based competition between prefrontal and dorsolateral striatal systems for behavioral control. Nat Neurosci 8, 1704–1711 (2005).

44. Silvetti, M., Vassena, E., Abrahamse, E. & Verguts, T. Dorsal anterior cingulate-brainstem ensemble as a reinforcement meta-learner. PLoS Comput Biol. 14, e1006370 (2018).

45. Pearce, J.M. & Mackintosh, N.J. Two theories of attention: a review and a possible integration. (Oxford University Press, New York; 2010).

46. Suzuki, M. & Gottlieb, J. Distinct neural mechanisms of distractor suppression in the frontal and parietal lobe. Nature Neuroscience 16, 98–104 (2013).

47. Gersch, T.M., Foley, N.C., Eisenberg, I. & Gottlieb, J. Neural correlates of temporal credit assignment in the parietal lobe. PLoS One 9, e88725 (2014).

48. Foley, N.C., Kelley, S.P., Mhatre, H., Lopes, M. & Gottlieb, J. Parietal neurons encode expected gains in instrumental information. Proceedings of the National Academy of Science 114, E3315–E3323 (2017).

49. Oostenveld, R., Fries, P., Maris, E. & Schoffelen, J.-M. FieldTrip: Open source software for advanced analysis of MEG, EEG, and invasive electrophysiological data. Comput. Intell. Neurosci. 156869 (2011).

